# Curcumin Analogues as the Inhibitors of TLR4 Pathway in Inflammation and Their Drug Like Potentialities: A Computer-based Study

**DOI:** 10.1101/2020.01.27.921528

**Authors:** Md. Asad Ullah, Fatema Tuz Johora, Bishajit Sarkar, Yusha Araf, MD. Hasanur Rahman

## Abstract

In this study Curcumin and their different analogues have been analyzed as the inhibitors of signaling proteins i.e., Cycloxygenase-2 (COX-2), Inhibitor of Kappaβ Kinase (IKK) and TANK binding kinase-1 (TBK-1) of Toll Like Receptor 4 (TLR4) pathway involved in inflammation using computational tools. Multiple analogues showed better binding affinity than the approved drugs for the respective targets. Upon continuous computational exploration 6-Gingerol, Yakuchinone A and Yakuchinone B were identified as the best inhibitors of COX-2, IKK and TBK-1 respectively. Then their drug like potentialities were analyzed in different experiments where they also performed sound and similar. Hopefully, this study will uphold the efforts of researchers to identify anti-inflammatory drugs from natural sources.

## 1. Introduction

**Inflammation** is delineated as normal biological as well as immune response against harmful stimuli including pathogens (bacteria, virus), toxins, stress, radiation, damaged cells etc. It is one of the protective mechanism of an organism to vanish invading stimulation, wound healing and for the restoration body’s normal physiology [1][2]. Inflammation is the result of several biological processes. Tissue injury or infection triggers inflammation as well as subsequent inflammatory cascade. Typical inflammatory cascade consists of four components (**Figure 1**).

**Figure 1:**
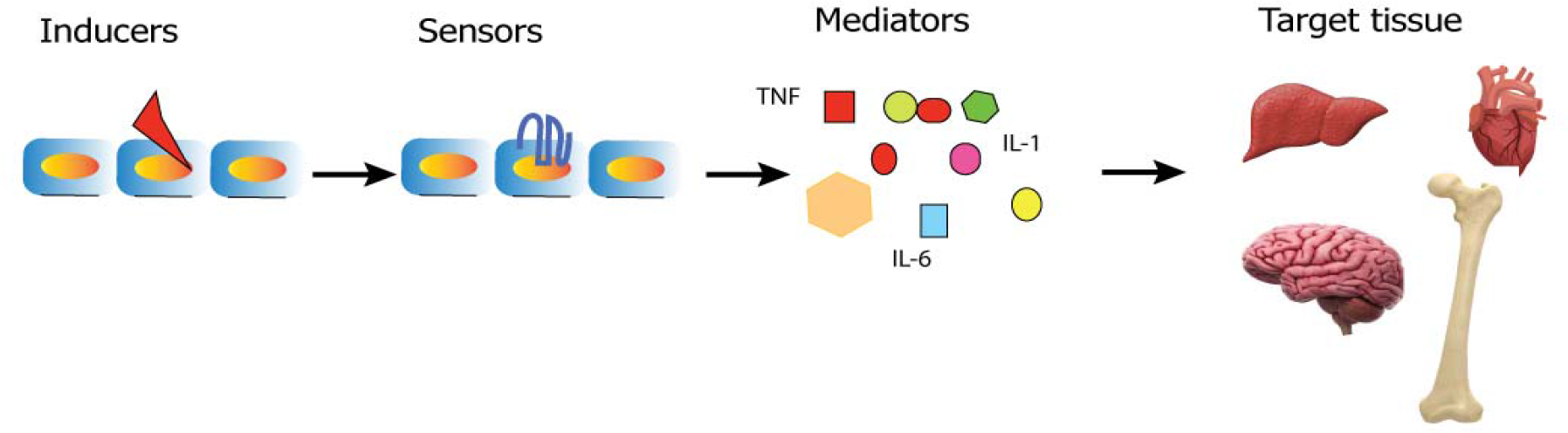
Components of inflammatory cascade. Inducers activate the sensors that results in the release of inflammatory mediators and then inflammatory mediators carry out the events that result in inflammation in target tissue.

(i). **Inducers**: Exogenous immune inducers (pathogens, virus, bacteria, allergens etc.) or endogenous inducers (damaged cells, stress) results in infection. (ii). **Sensors:** Specific receptors i.e., toll like receptors (sensors) recognize these inducers. These receptors are localized on the immune cells (mast cells, dendritic cells, macrophage). (iii)**. Mediators**: After that immune cells release different types of mediators (TNF, IL-1, IL-6). Mediators vary depending on the inducers type. (iv). **Inflammation in Target Tissue**: Immune mediators elicit their effects (dilation of blood vessels, increased vascular permeability, movement of leukocytes from blood vessels to inured area etc.) on target tissues [3][4].

There are two forms of inflammation i.e., acute inflammation and chronic inflammation. Acute inflammation is pictured as immediate, short-term response and innate immunity to injury. Augmented movement of leukocytes from blood to the injured area results in acute inflammation within minutes or hours and lasted for short time to irradiate harmful stimuli. When the inflammatory responses last for long periods, it leads to chronic inflammation. Chronic inflammation leads to more complicated physiological condition and several diseases[5] i.e., neuroinflammation (brain), metabolic disorders (liver/ pancreas) [6], osteoporosis (bone) [7][8], cardiovascular disease (heart)[9], cancer [10], rheumatoid arthritis [11], obesity, asthma, Alzheimer’s disease [12]etc.

### 1. 2. Role of Toll Like Receptor 4 (TLR4) Pathway in Inflammation

In response to inflammatory inducers (signals i.e., bacterial lipopolysaccharides) pattern recognition receptors as well as transmembrane receptors play a pivotal role in initiation of immune response, among these, Toll like receptors are most significant [13]. TLR4 senses the harmful stimuli and recruits the coordinate activation of two distinct transcription factors i.e., Nuclear factor kappaβ (NF-кB) and Interferon Regulatory Factor 3 (IRF3) (**Figure 2**) [14]-[16].

**Figure 2:**
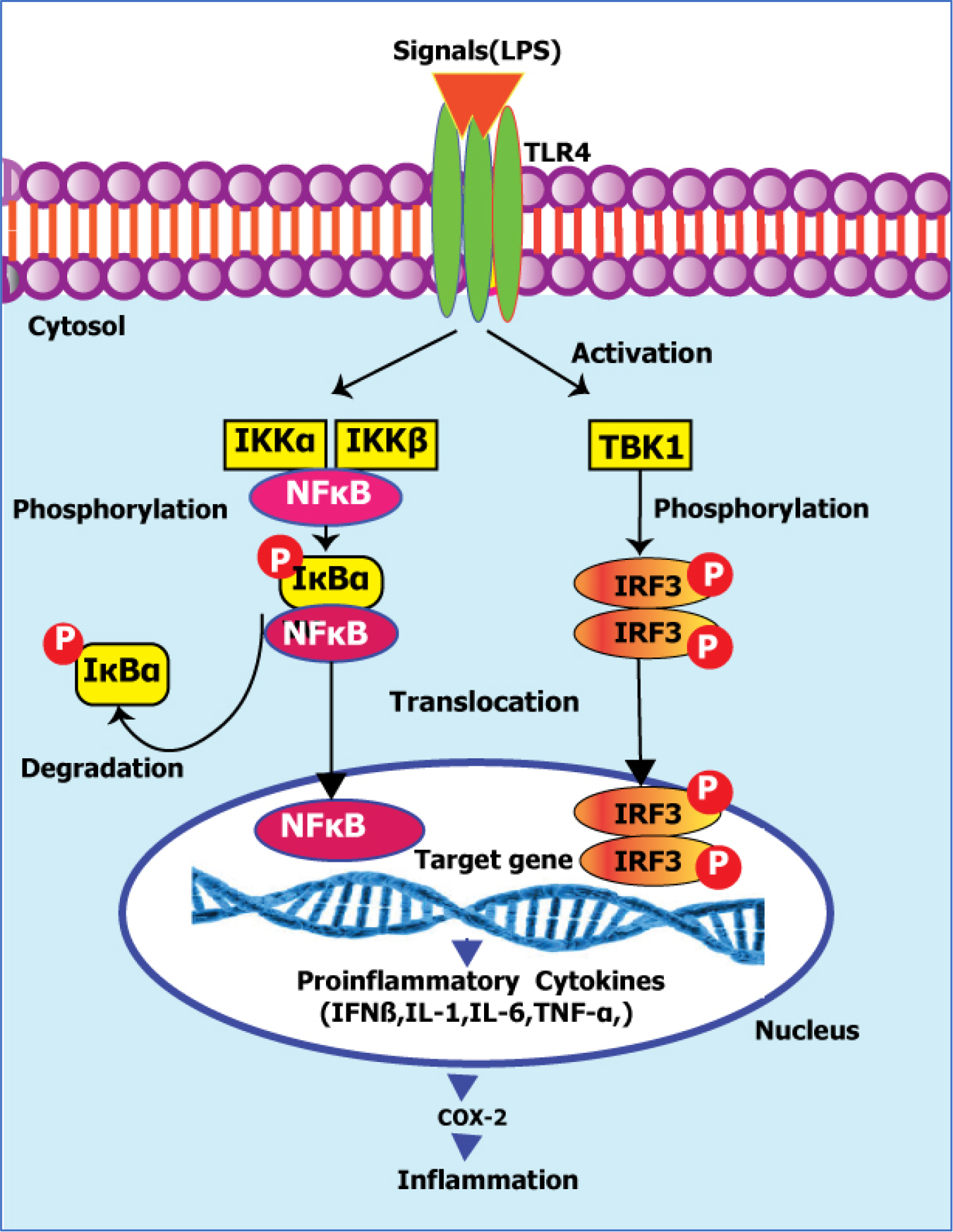
TLRs pathway. Sensing harmful stimuli, Toll like receptor4 (TLR4) gets activated and recruits the coordinate activation of two distinct transcription factors i.e., NF-кB and IRF3. Activated TLR4 at first activates both canonical (IKKα/IKKβ) and noncanonical (TBK1) IkappaB kinases (IКB) and these kinases activate the NF-кB and IRF3 transcription factor respectively which leads to their translocation into nucleus and facilitates the transcription of proinflammatory genes.

In the resting cells, NF-кB is isolated by inhibitory proteins called, IkappaB/IКB kinase (IKK) in cytoplasm where these proteins act like a mask and inactivate NF-кB. In response to signals TLR4 is activated which in turn activate both canonical IкB kinases (IкB kinase α/β-IKKα, IKKβ) and noncanonical IкB kinase, i.e., TBK1 (TANK binding kinase-1)[17]. Activated IKKα/IKKβ (IKK complex) phosphorylate IкBα which is ubiquitinated and degraded and activate NF-кB. Activated TBK1 results in phosphorylation of IRF3 which in turn leads to formation of dimer complex. Both the activated NF-кB and IRF3 dimer translocate in the nucleus and facilitate the transcription of the gene of proinflammatory cytokines (TNF-α, IL-1, IL-6, IFN-β). These proinflammatory cytokines cause fever, inflammation, pain, tissue destruction in human body. NF-кB plays a substantial role in the regulation and production of COX-2 which acts on arachidonic acid and facilitates it conversion into prostaglandins [18]-[20]. COX-2 expression is inducible and mainly expressed in immune cells and elicits pathophysiological functions. COX-2 produces high level PGs which play a substantial role in the inflammatory response where they act as inflammatory mediators. Dramatically augmented level of PGs develops cardinal signs of inflammation as well as pain, swelling and other physiological disorders [21].

### 1.3. Current treatments of inflammation and limitation

Nonsteroidal anti-inflammatory drugs (NSAIDs) are most commonly prescribed drugs for inflammation. These drugs reduce pain and immune response by inhibiting COX enzymes, TBK1 (Amlexanox, FDA approved drug [22]) and IKK kinases (Quilonoxaline) that take part in the synthesis of prostaglandins[23][24]. These drugs are classified into two groups i.e., Non-selective NSAIDs (aspirin, ibuprofen, naproxen etc.) and selective NSAIDs (celecoxib, rofecoxib). [25]-[29]. Treatment of inflammation with selective NSAIDs is lucrative. Because selective NSAIDs, selectively inhibits COX-2. As a result, body’s homostatic functions are carried out perfectly. Recent studies show that, selective NSAIDs cause cardiovascular complications, disturbs normal renal function as COX-2 is involved in renal development [30][31]. As the use of these existing drugs is very challenging, plant-based medicines are more desirable as they found to have less side effects [32].

### 1.4. Curcumin Analogues as Anti-inflammatory Agents

Plants are sources of wide varieties of phytochemicals which provide range of therapeutic benefits in human body [33][34]. Curcumin is a dietary yellow pigment as well as potent anti-inflammatory agent, found in turmeric (*Curcuma longa)* as it inhibits the prostaglandin (PG) synthesis which is crucial for the initiation of inflammation. There are various types of analogues of Curcumin, found in other plants of the mother nature [35]. Curcumin has been proven to have anti-inflammatory activities and down regulate the NF-кB activation and TLR4 pathway activity in inflammation in many pathways [36]-[38]. In a laboratory experiment Curcumin has been shown to inhibit COX-2 as well as COX-1 activity by 50% in a concentration of 15 µM with slight selectivity [39]. In yet other studies Curcumin has been shown to inhibit IKK activity and downregulate NF-кB activation [40]. Curcumin Analogues Yakuchinone A and Yakuchinone B were also reported to inhibit COX-2 activity in laboratory experiment [41]. 6-Shogaol has similar structure as of Curcumin and a structural analogue of 6-Gingerol is an inhibitor of TBK-1[42]. In this experiment different Curcumin analogues have been analyzed to understand their inhibitory effects on multiple signaling proteins (targets) involved in inflammation based on the hypothesis that since they have similar structure and few analogues have target specific anti-inflammatory activity so one or more analogue(s) could have even better activity.

## 2. Materials and Methods

A total of 14 compounds comprising Curcumin, its derivatives and analogs were selected from literature review. Then they were subjected to drug likeness property analysis, molecular docking, and other experiments to identify best inhibitors of the respective targets (**Figure 3**).

**Figure 3:**
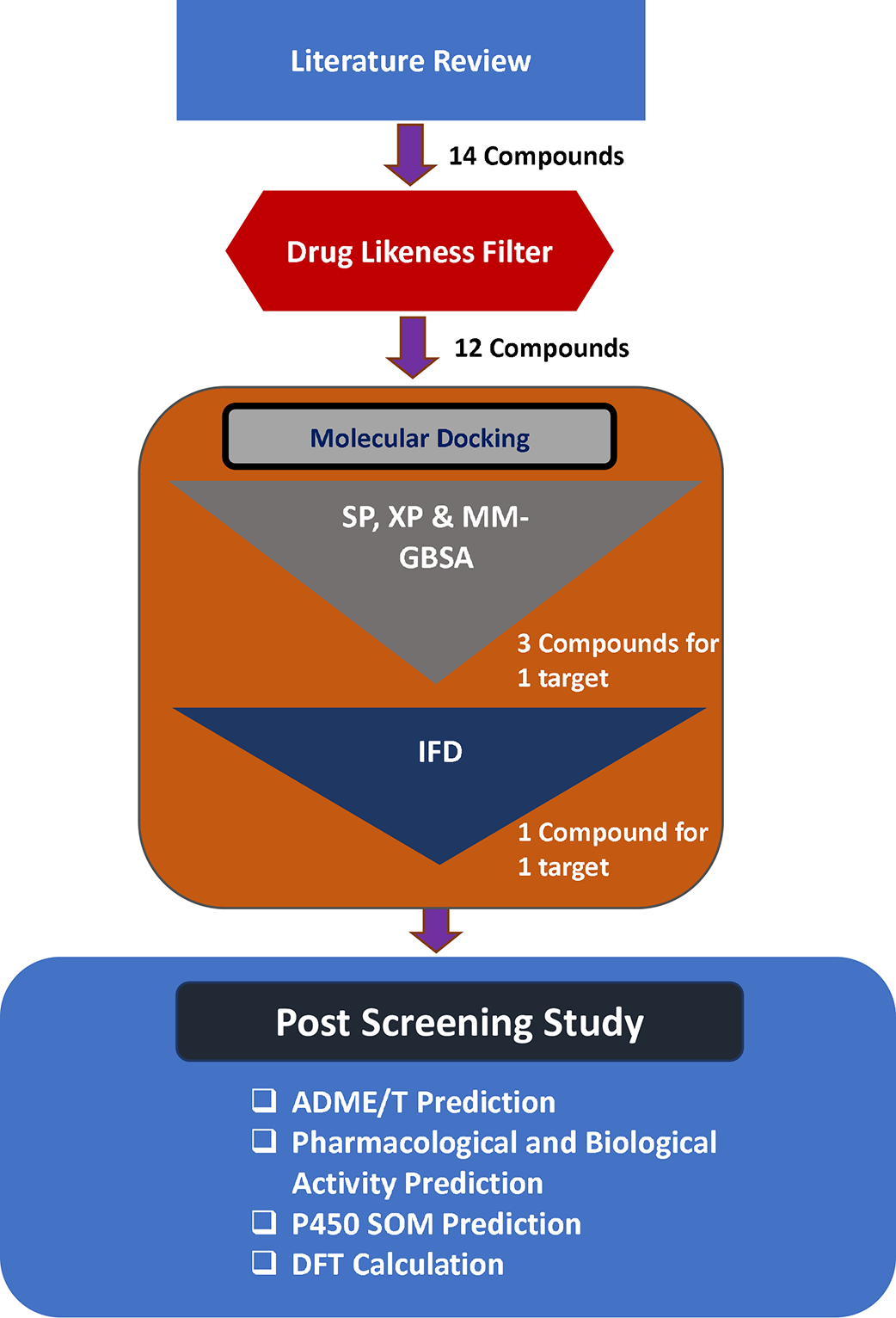
Strategies employed in the overall study.

### 2.1. Drug Likeness Property Analysis

The selected ligand molecules were analyzed to determine whether they obey Lipinski’s rule of five or not which states that a drug is considered to have poor bioavailability and low permeation which violates the standard rule [55][56]. Canonical smile of each intended ligand molecule was retrieved from PubChem database (https://pubchem.ncbi.nlm.nih.gov/) and was then analyzed using the Molonspiration Cheminformatics server (https://www.molinspiration.com/cgi-bin/properties) for different drug-likeness parameters (**Table 3**)[57][58]. Compounds violating the rule were then opted out from consideration and further evaluation.

**Table 1:** Curcumin and it’s natural derivatives from turmeric.

| Compound name | Source | PubChem CID | Reference |
| --- | --- | --- | --- |
| Bisdemethoxycurcumin | Turmeric<br>(*Curcuma longa*) | 5315472 | [43] |
| Curcumin | Turmeric<br>(*Curcuma longa*) | 969516 | [44] |
| Cyclocurcumin | Turmeric<br>(*Curcuma longa*) | 69879809 | [45] |
| Demethoxycurcumin | Turmeric<br>(*Curcuma longa*) | 5469424 | [43] |

**Table 2:** Curcumin analogs from mother nature.

| Compound name | Source | PubChem CID | Reference |
| --- | --- | --- | --- |
| 6-Gingerol | Ginger<br>(*Zingiber officinale Roscoe*) | 442793 | [46] |
| 6-Paradol | Ginger<br>(*Zingiber officinale Roscoe*) | 94378 | [47][48] |
| 6-Shogaol | Ginger<br>(*Zingiber officinale*) | 5281794 | [49] |
| Cassumunin A | Ginger<br>(*Zingiber cassumunar*) | 10460395 | [50] |
| Cassumunin B | Ginger<br>(*Zingiber cassumunar*) | 10054109 | [50] |
| Dehydrozingerone | Ginger (*Zingiber officinale Roscoe*) | 5354238 | [51] |
| Dibenzoylmethane | Licorice<br>(*Glycyrrhiza echinata*) | 8433 | [52] |
| Isoeugenol | Cloves<br>(*Eugenia caryophyllus*) | 853433 | [53] |
| Yakuchinone A | Galanga<br>(*Alpinia officinarum*) | 133145 | [54] |
| Yakuchinone B | Galanga<br>(*Alpinia officinarum*) | 6440365 | [54] |

**Table 3:** Results of drug likeness property analysis. HBA; Hydrogen bond acceptors; HBD: Hydrogen bond donors, nROTB: Number of rotatable bonds; TPSA: Topological Polar Surface Area.

| SL. No. | Compound Name | MW (g/mol) | miLogP | HBA | HBD | nROTB | TPSA (Å<sup>2</sup>) | Lipinski Violation |
| --- | --- | --- | --- | --- | --- | --- | --- | --- |
| **Rule** |  | <500 | ≤5 | <10 | <5 | ≤10 |  |  |
| 01. | 6-Gingerol | 294.39 | 3.22 | 4 | 2 | 10 | 66.76 | 0 |
| 02. | 6-Paradol | 278.39 | 4.60 | 3 | 1 | 10 | 46.53 | 0 |
|  | 6-Shogaol | 276.38 | 4.35 | 3 | 1 | 9 | 46.53 | 0 |
| 03. | Bisdemethoxycurcumin | 308.33 | 2.67 | 4 | 2 | 6 | 74.60 | 0 |
| 04. | Cassumunin A | 558.63 | 4.96 | 8 | 2 | 13 | 111.53 | 1 |
| 05. | Cassumunin B | 588.65 | 4.77 | 9 | 2 | 14 | 120.77 | 1 |
| 06. | Curcumin | 368.38 | 2.30 | 6 | 2 | 8 | 93.07 | 0 |
| 07. | Cyclocurcumin | 368.38 | 3.03 | 6 | 2 | 5 | 85.23 | 0 |
| 08. | Dehydrozingerone | 192.21 | 1.55 | 3 | 1 | 3 | 46.53 | 0 |
| 09. | Demethoxycurcumin | 338.36 | 2.48 | 5 | 2 | 7 | 83.83 | 0 |
| 10. | Dibenzoylmethane | 224.26 | 2.88 | 2 | 0 | 4 | 34.14 | 0 |
| 11. | Isoeugenol | 164.20 | 2.38 | 2 | 1 | 2 | 29.46 | 0 |
| 12. | Yakuchinone A | 312.41 | 4.24 | 3 | 1 | 9 | 46.53 | 0 |
| 13. | Yakuchinone B | 310.39 | 4.27 | 3 | 1 | 8 | 46.53 | 0 |

### 2.2. Molecular Docking Experiment

#### 2.2.1. Protein Preparation

Three dimensional crystallized structure of Human COX-2 (PDB ID:5F1A), Inhibitor of kappaB kinase beta (PDB ID: 3RZF) and Tank Binding Kinase-1 (PDB ID:5W5V) were downloaded in PDB format from Protein Data Bank (www.rcsb.org) [59]-[61]. The structures were then prepared and processed using the Protein Preparation Wizard in Maestro Schrödinger Suite (v11.4). Bond orders were assigned to the structures, hydrogens were added to heavy atoms. All of the water molecules were erased from the atoms, missing side chains were added to the protein backbone using Prime and het states were generated with Epik at pH 7 ± 2 [62]. At last, the structures were refined and then minimized utilizing Optimized Potentials for Liquid Simulations force field (OPLS_2005). Minimization was performed setting the greatest substantial particle RMSD (root-mean-square-deviation) to 30 Å and any extraordinary water under 3H-bonds to non-water was again eradicated during the minimization step.

#### 2.2.2. Ligand Preparation

A total of 12 ligand molecules except those that violated Lipinski’s rule of five were downloaded in sdf format from PubChem database (https://pubchem.ncbi.nlm.nih.gov/) [63]. These structures were then processed and prepared using the LigPrep wizard of Maestro Schrödinger suite [64]. Minimized 3D structures of ligands were generated using Epik2.2 and within pH 7.0 +/-2.0 in the suite. Minimization was again carried out using OPLS_2005 force field which generated maximum 32 possible stereoisomers depending on available chiral centers on each molecule.

#### 2.2.3. Receptor Grid Generation

Grid usually confines the active site to specific area of the receptor protein for the ligand to dock specifically within selected area. Receptor grid was generated using default Van der Waals radius scaling factor 1.0 and charge cutoff 0.25 which was then subjected to OPLS_2005 force field for the minimized structure in Glide [65]. A cubic box was then generated around the active site (co-crystallized reference ligand) of the target molecules. The grid box dimension was then adjusted to 14 Å ×14 Å ×14 Å for docking to be carried out.

#### 2.2.4. Glide Standard Precision (SP) and Extra Precision Ligand Docking

Extra precision (XP) ligand docking is more accurate for small number of ligand molecules than standard precision (SP) ligand docking which is recommended for large compound libraries [66]. But both of the docking methods were applied for the selected ligand molecules and intended targets for making comparison. The Van der Waals radius scaling factor and charge cutoff were set to 0.80 and 0.15 respectively for all the ligand molecules under study. Final score was assigned according to the pose of docked ligand within the binding cleft of the receptor molecules. Best possible poses and types of ligand-receptor interactions were then analyzed utilizing Discovery Studio Visualizer (v4.5) (**Figure 4**) [67].

**Figure 4:**
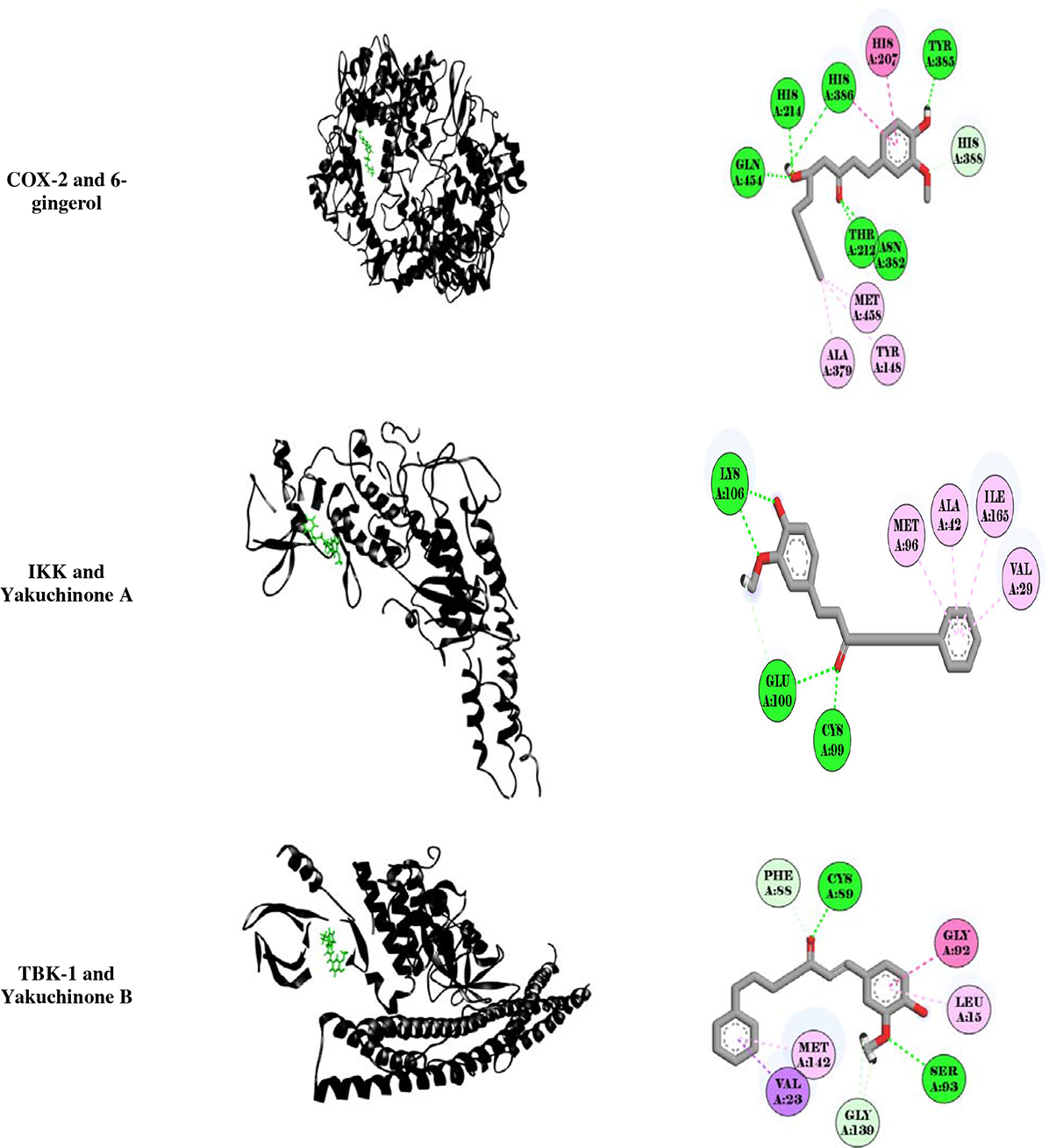
(A) Three dimensional representation of best possible poses of ligand molecules (green, stick) inside the binding pocket of intended target (black, ribbon). (B) Two dimensional representation of ligand-receptor interaction. Interacting amino acids are represented in three letter code and their respective number in specific chain inside disc. Dotted lines represent type of interactions: Green: Conventional hydrogen bond; Light green: Pi-Donor and Carbon hydrogen Bond. Pink: Amide-Pi stacked, Pi-Pi stacked and Pi-Pi T shaped; Light pink: Alkyl and Pi-Alkyl.

#### 2.2.5. Prime MM-GBSA Rescoring

After SP and XP ligand docking the ligands were then again subjected to Molecular mechanics – generalized born and surface area (MM-GBSA) rescoring with the help of Prime module of Maestro Schrödinger suite for further evaluation. This technique utilizes an implicit solvent which then assigns more accurate scoring function that then improves the overall free binding affinity score upon the reprocessing of the docked complex [66][68]. It combines OPLS molecular mechanics energies (E_MM_), surface generalized born solvation model for polar solvation (G_SGB_), and a nonpolar salvation term (G_NP_) for total free energy (ΔG_bind_) calculation. The total free energy of binding was calculated by the following equation:
**ΔG_bind_=G_complex_-(G_protein_-G_ligand), where, G=E_MM_+G_SGB_+G_NP__**

The result of SP docking, XP docking and MM-GBSA rescoring is summarized in **Table 4**.

**Table 4:** Results of SP, XP docking and Free binding energy calculation between intended target and ligand molecules.

| Receptor Name | Compound Name | SP Docking Score (Kcal/mol) | XP Docking Score (Kcal/mol) | Glide Energy | Glide-ligand Efficiency | MM-GBSA $\Delta G_{\text{bind}}$ (Kcal/mol) |
| --- | --- | --- | --- | --- | --- | --- |
| COX-2 | Celecoxib (control) | -4.854 | -6.139 | -29.387 | -0.186 | -39.873 |
|  | Cyclocurcumin | -7.394 | -7.962 | -49.949 | -0.336 | -46.310 |
|  | 6-Gingerol | -5.368 | -6.693 | -52.051 | -0.271 | -57.913 |
|  | Demethoxycurcumin | -5.909 | -6.622 | -50.027 | -0.225 | -55.740 |
| IKK | Salfasalazine (control) | -4.949 | -5.131 | -43.935 | -0.183 | -55.510 |
|  | Yakuchinone A | -6.283 | -7.791 | -38.429 | -0.339 | -69.953 |
|  | Yakuchinone B | -6.918 | -6.895 | -36.961 | -0.300 | -68.737 |
|  | 6-Paradol | -4.494 | -5.555 | -34.138 | -0.278 | -68.553 |
| TBK-1 | Fostamatinib (control) | -6.980 | -5.518 | -63.716 | -0.138 | -53.750 |
|  | Yakuchinone B | -7.066 | -7.837 | -38.540 | -0.341 | -56.043 |
|  | Yakuchinone A | -7.189 | -7.393 | -38.764 | -0.321 | -52.469 |
|  | Curcumin | -6.273 | -6.266 | -43.968 | -0.232 | -52.819 |

#### 2.2.6. Induced Fit Docking

Three compounds were selected based on the lowest MM-GBSA score for each of the receptor molecule which were then used for further evaluation since it is more robust scoring method. At this stage, different scores of three best docked compounds were compared with one approved known inhibitor (control) i.e., Celecoxib, Salfasalazine and Fostamatinib for COX-2, IKK and TBK-1 (**Table 5**) receptor respectively [69]. After that the best three ligands for each receptor were subjected to induced fit docking (IFD) which is even more accurate docking method to generate the native poses of the ligands [70]. Again OPLS_2005 force field was applied after generating grid around the co-crystallized ligand of the receptor and this time the best five ligands were docked rigidly. Receptor and Ligand Van Der Waals screening was set at 0.70 and 0.50 respectively, residues within 2 Å were refined to generate 2 best possible posses with extra precision. Best performing ligand was selected according to the IFD score for one receptor molecule (**Table 6**). Then all three selected ligands for three receptors were used in the next phases of this study.

**Table 5:** Result of induced fit docking (IFD) between best performing ligand and respective target.

| Target Name | Ligand Name | IFD Score (Kcal/mol) | XP Gscore (Kcal/mol) | Interacting Amino Acids | Bond Distance (Å) | Type of Interaction | Interaction Category |
| --- | --- | --- | --- | --- | --- | --- | --- |
| COX-2 | 6-Gingerol | -2474.460 | -9.264 | Gln454 | 2.01 | Hydrogen bond | Conventional |
|  |  |  |  | His386 | 4.74 | Pi-Pi Stacked | Hydrophobic |
|  |  |  |  | HIS207 | 4.48 | Pi-Pi T Shaped | Hydrophobic |
|  |  |  |  | His214 | 2.29 | Hydrogen bond | Conventional |
|  |  |  |  | Tyr385 | 1.93 | Hydrogen bond | Conventional |
|  |  |  |  | Thr212 | 3.60 | Hydrogen bond | Conventional |
|  |  |  |  | His388 | 2.03 | Hydrogen bond | Non-conventional |
|  |  |  |  | Asn382 | 2.52 | Hydrogen bond | Conventional |
|  |  |  |  | Met458 | 4.84 | Pi-Alkyl | Hydrophobic |
|  |  |  |  | Tyr148 | 4.15 | Pi-Alkyl | Hydrophobic |
|  |  |  |  | Ala379 | 3.71 | Pi-Alkyl | Hydrophobic |
| IKK | Yakuchinone A | -1081.370 | -9.111 | Lys106 | 5.46 | Hydrogen bond | Conventional |
|  |  |  |  | Met96 | 4.26 | Pi-Alkyl | Hydrophobic |
|  |  |  |  | Lys106 | 2.00 | Hydrogen bond | Conventional |
|  |  |  |  | Ala42 | 4.27 | Pi-Alkyl | Hydrophobic |
|  |  |  |  | Glu100 | 2.72 | Hydrogen bond | Conventional |
|  |  |  |  | Ile165 | 5.19 | Pi-Alkyl | Hydrophobic |
|  |  |  |  | Glu100 | 3.04 | Hydrogen bond | Non-conventional |
|  |  |  |  | Val29 | 3.93 | Pi-Alkyl | Hydrophobic |
|  |  |  |  | Cys99 | 1.96 | Hydrogen bond | Conventional |
| TBK-1 | Yakuchinone B | -1309.340 | -7.971 | Val23 | 2.71 | Pi-Sigma | Hydrophobic |
|  |  |  |  | Met142 | 5.21 | Pi-Alkyl | Hydrophobic |
|  |  |  |  | Gly139 | 2.60 | Hydrogen bond | Non-conventional |
|  |  |  |  | Ser93 | 2.84 | Hydrogen bond | Conventional |
|  |  |  |  | Leu15 | 5.17 | Pi-Alkyl | Hydrophobic |
|  |  |  |  | Gly92 | 4.68 | Amide-Pi Stacked |  |
|  |  |  |  | Cys89 | 2.09 | Hydrogen bond | Conventional |
|  |  |  |  | Phe88 | 2.76 | Hydrogen bond | Non-conventional |
|  |  |  |  | Gly139 | 2.81 | Hydrogen bond | Non-conventional |

**Table 6:**
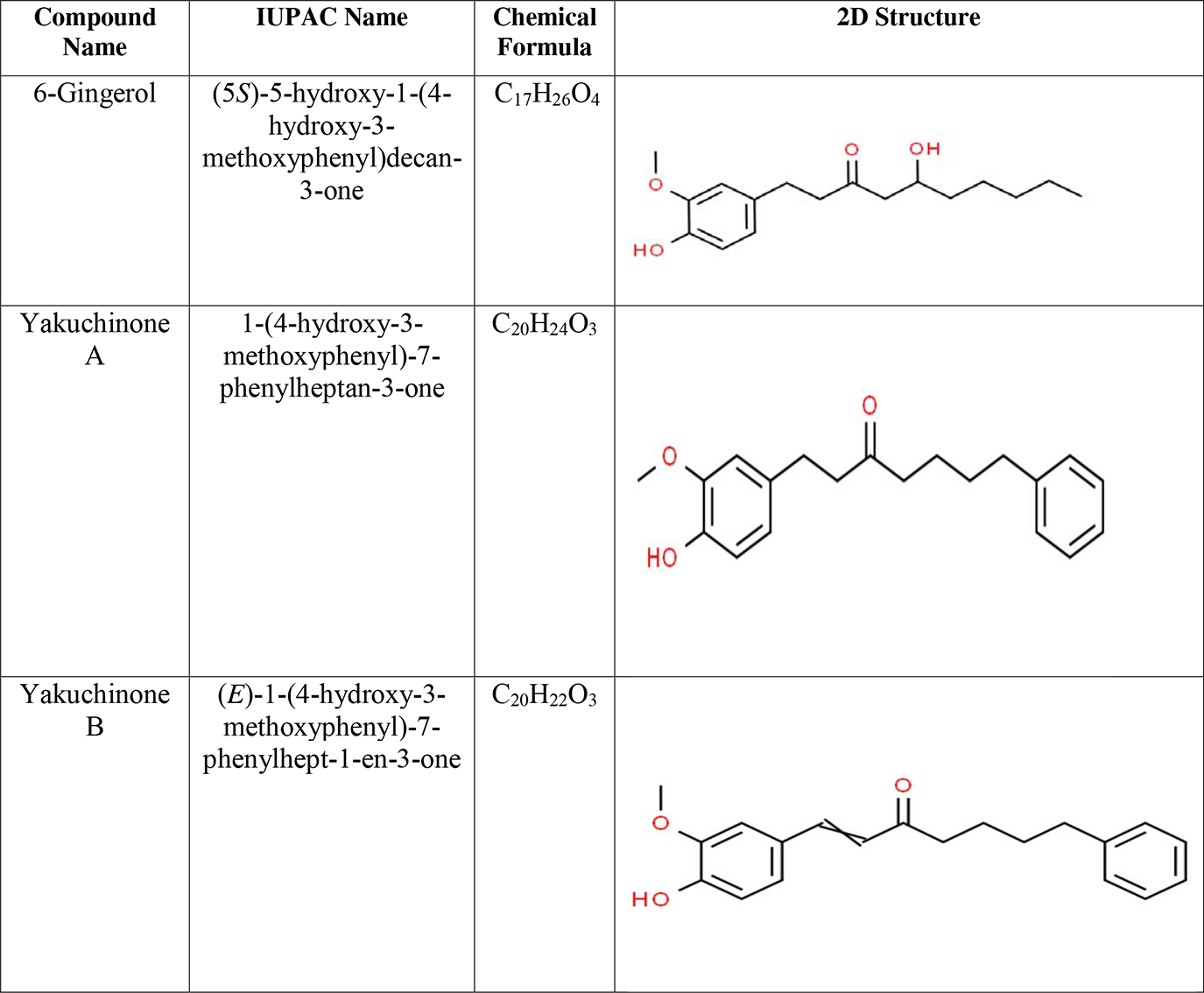
Best performed ligand in overall docking experiment.

| Compound Name | IUPAC Name | Chemical Formula | 2D Structure |
| --- | --- | --- | --- |
| 6-Gingerol | (5*S*)-5-hydroxy-1-(4-hydroxy-3-methoxyphenyl)decan-3-one | C<sub>17</sub>H<sub>26</sub>O<sub>4</sub> |  |
| Yakuchinone A | 1-(4-hydroxy-3-methoxyphenyl)-7-phenylheptan-3-one | C<sub>20</sub>H<sub>24</sub>O<sub>3</sub> |  |
| Yakuchinone B | (*E*)-1-(4-hydroxy-3-methoxyphenyl)-7-phenylhept-1-en-3-one | C<sub>20</sub>H<sub>22</sub>O<sub>3</sub> |  |

### 2.3. Ligand-based ADME/T Prediction

*In silico* prediction of ADME/T profile of candidate drug molecule helps to increase the success rate of drug discovery expenditure [71][72]. Canonical smiles of the best three ligand were used to predict drug like potential and tentative pharmacokinetic and pharmacodynamic parameters. ADME/T profile of each ligand was predicted using admetSAR 2.0 (http://lmmd.ecust.edu.cn/admetsar2/) and pkCSM server (http://biosig.unimelb.edu.au/pkcsm/) [73][74] (**Table 7**).

**Table 7:**
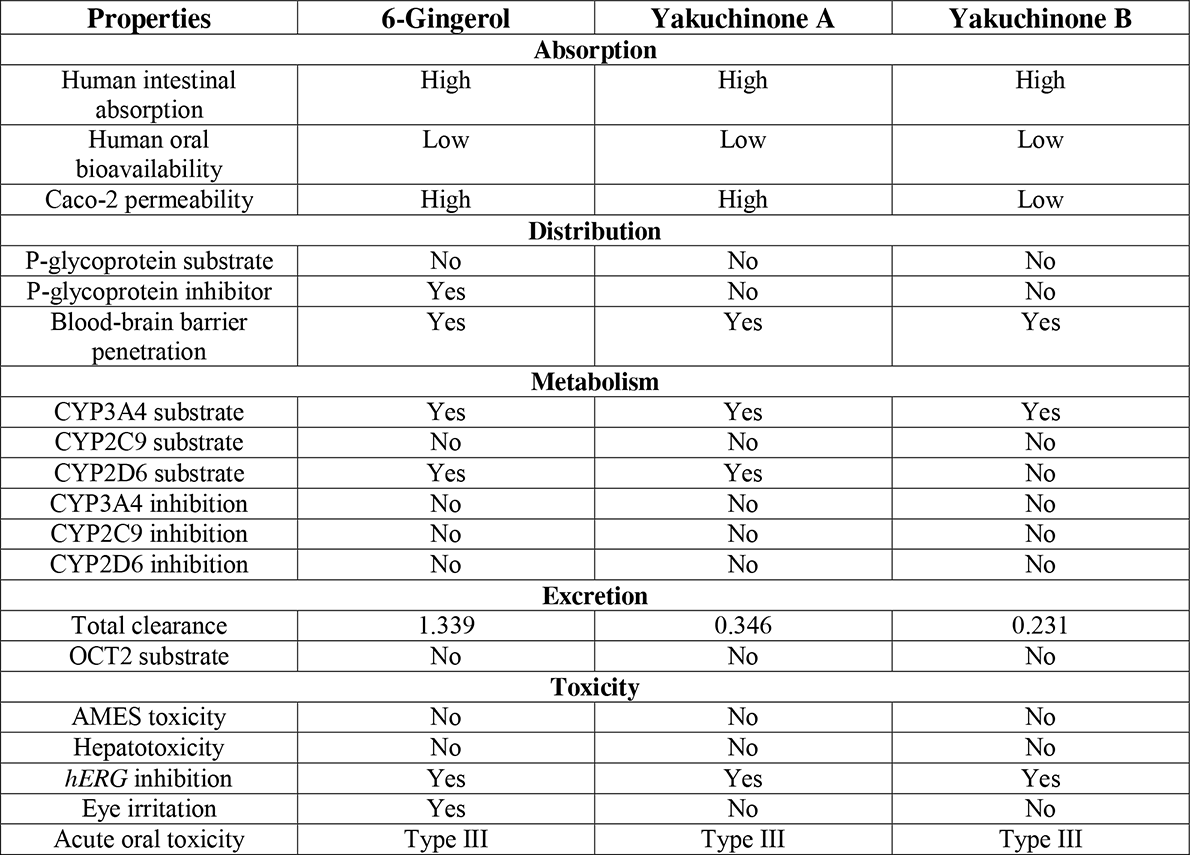
Results of ADME/T tests of best selected ligands. OCT2: Organic Cation Transporter 2; *hERG:* Human ether-a-go-go related gene, CYP: Cytochrome P450

### 2.4. Pharmacological and Biological Activity Prediction

Pharmacological and biological activities of the best ligand molecules were predicted using PASS (Prediction of Activity Spectra of Substances) Online (http://www.pharmaexpert.ru/passonline/) and Molinspiration Cheminformatics servers respectively (**Table 8** & **9**) [58][75]. These tools predict the tentative activities of compounds based on structure activity relationship (SAR) in correlation with a known compound existing in the database.

**Table 8:** Result of Pharmacological Activity prediction of selected ligand molecules. Pa>0.7: Compound is very likely to have activity; Pa>0.5: Compound is likely to have activity; Pa>0.3: Compound is less likely to have activity.

| Pharmacological Activity | 6-Gingerol |  | Yakuchinone A |  | Yakuchinone B |  |
| --- | --- | --- | --- | --- | --- | --- |
|  | Pa | Pi | Pa | Pi | Pa | Pi |
| Antiinflammatory | 0.532 | 0.532 | 0.497 | 0.058 | 0.576 | 0.037 |
| Antiinflammatory, intestinal | 0.566 | 0.004 | 0.437 | 0.015 | 0.519 | 0.006 |
| Antiinflammatory, ophthalmic | 0.343 | 0.343 | 0.330 | 0.043 | 0.329 | 0.044 |
| MAP kinase stimulant | 0.482 | 0.049 | 0.667 | 0.008 | 0.658 | 0.009 |
| TNF expression inhibitor | 0.633 | 0.010 | 0.595 | 0.014 | 0.804 | 0.004 |
| JAK2 expression inhibitor | 0.679 | 0.020 | 0.863 | 0.004 | 0.945 | 0.002 |
| NF kappa A inhibitor | 0.228 | 0.115 | 0.278 | 0.048 | 0.266 | 0.059 |
| NF kappa B inhibitor | 0.264 | 0.015 | 0.240 | 0.018 | 0.341 | 0.009 |
| Macrophage colony stimulating factor agonist | 0.762 | 0.007 | 0.687 | 0.015 | 0.569 | 0.040 |
| Cyclooxygenase substrate | 0.326 | 0.011 | 0.230 | 0.058 | 0.224 | 0.064 |

### 2.5. P450 Site of Metabolism Prediction

*In silico* analysis of potential sites of metabolism of candidate drug metabolism provides insights into the metabolic vulnerability of the molecule inside in human body which then drives the *in vitro* assay [76]. Best metabolism sites of best three ligands to three isoforms i.e. CYP3A4, CYP2D6 and CYP2C9 of Cytochrome P450 family of enzymes were predicted utilizing online-based RS-WebPredictor server (http://reccr.chem.rpi.edu/Software/RS-WebPredictor/) (**Figure 5**) [77].

**Figure 5:**
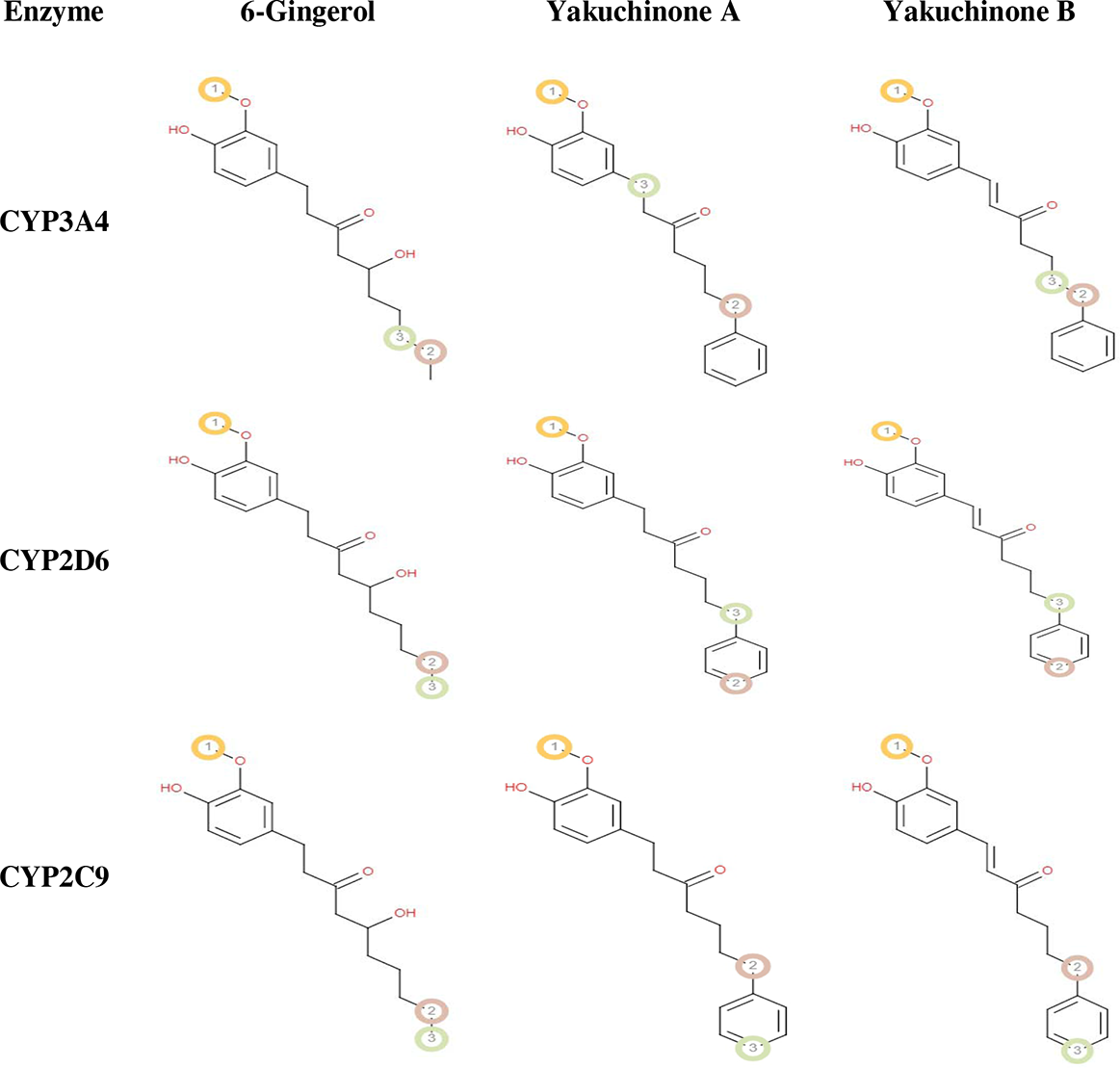
Results of P450 site of metabolism prediction. Best three vulnerable atoms are marked in encircled number.

### 2.6. DFT (Density Functional Theory) Calculation

Minimized ligand structures was obtained from LigPrep which were then used for DFT calculation using the Jaguar panel of Maestro Schrödinger Suite using Becke’s three-parameter exchange potential and Lee-Yang-Parr correlation functional (B3LYP) theory with 6-31G* basis set in the suite [78]-[81]. Quantum chemical properties i.e. surface properties (MO, density, potential) and Multipole moments were calculated along with HOMO (Highest Occupied Molecular Orbital) and LUMO (Lowest Unoccupied Molecular Orbital) energy. Global frontier orbital was analyzed along with hardness (**η**) and softness (**S**) of selected molecules were also calculated using the following equation as per Parr and Pearson interpretation and Koopmans theorem [82][83]. The result of DFT calculation is summarized in **Table 10**. HOMO and LUMO occupation of the ligands is illustrated in **Figure 6**.
**η=(HOMO□-LUMO□)/2, S=1/η**

**Figure 6:**
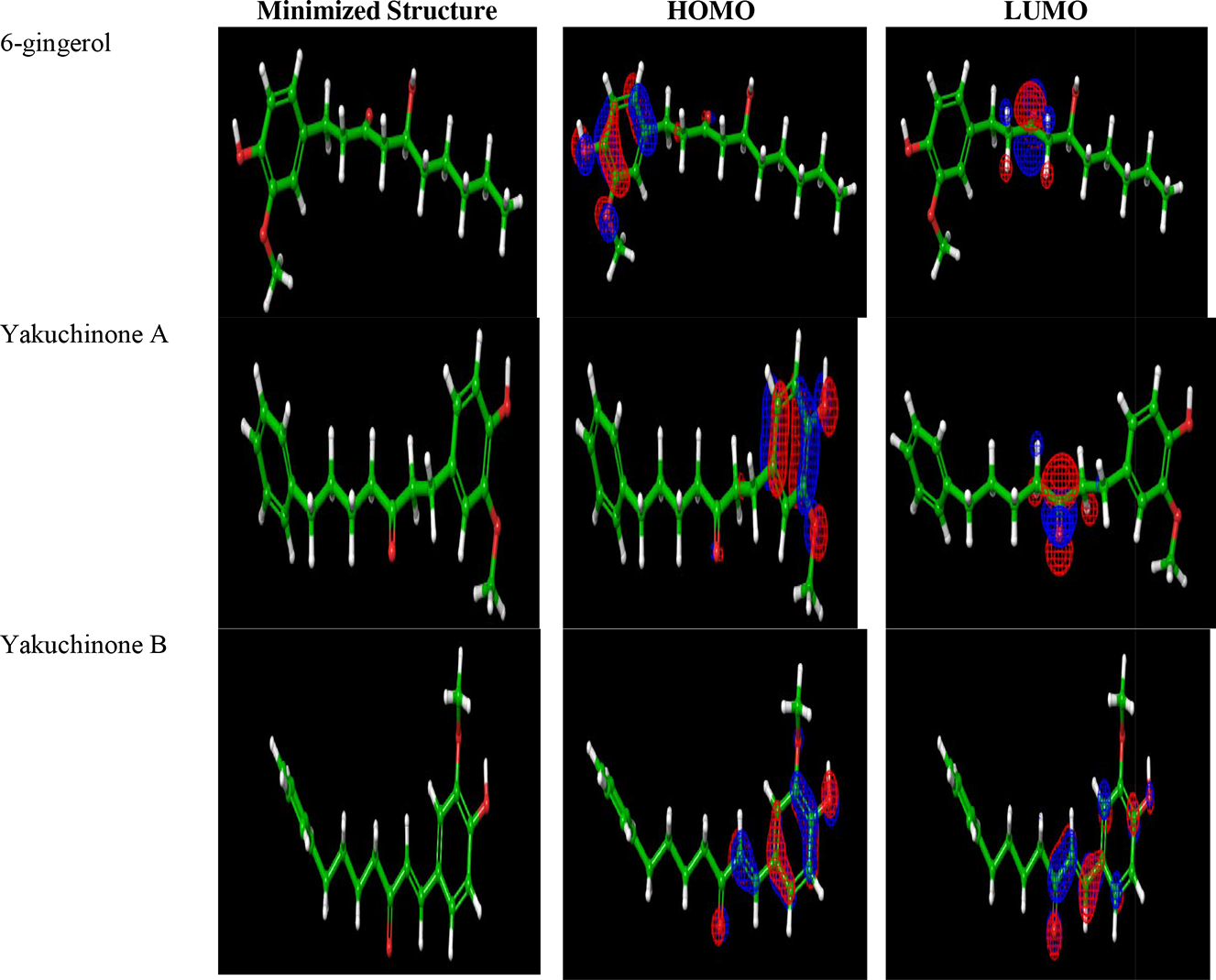
The HOMO and LUMO occupation for the selected compounds. Blue and red is positive and negative in its wave function.

## 3. Results

### 3.1. Drug Likeness Property

14 selected ligand molecules were analyzed to understand whether they comply Lipinski’s rule of five or not. Cassumunin A and Cassumunin B violated the standard rule and hence were removed for further consideration (**Table 3**). All other ligand molecules were reported to obey Lipinski’s rule of five. These 12 ligand molecules were then utilized in the next phases of the experiment.

Alongside the standard rule, the ligands were also analyzed for their Topological Polar Surface Area (TPSA), Isoeugenol was reported to have lowest TPSA and Curcumin was reported to have highest TPSA. Other ligands have TPSA within the moderate range among the highest and lowest values.

### 3.2. Molecular Docking Experiment

12 selected ligand molecules from the drug likeness property analysis were then utilized in the docking experiment against COX-2, IKK and TBK-1. Both standard precision (SP) and extra precision (XP) ligand docking were carried out. A slight variation between SP and XP ligand docking was observed. Then the docked complexes were utilized to calculate binding free energies and three best compounds with lowest binding energy were selected for each target molecule (**Table 4**). At this stage the docking parameters of approved known inhibitors (controls) of respective receptors were compared with that of the ligand of our specific interest.

Cyclocurcumin, 6-Gingerol and Demethoxycurcumin docked with COX-2 with −46.310 Kcal/mol, −57.913 Kcal/mol and −55.740 Kcal/mol free binding energies. And all of these ligand molecules showed lowest binding free energies than Celecoxib (−39.873 Kcal/mol) (control) (**Table 4**).

Yakuchinone A, Yakuchinone B and 6-Paradol docked with IKK with −69.953 Kcal/mol, −68.737 Kcal/mol and −68.553 Kcal/mol respectively. Control of this experiment (Salfasalazine) generated notably more positive free binding energy score (−55.510 Kcal/mol) than all compounds under investigation.

Again, Yakuchinone A, Yakuchinone B and Curcumin docked with TBK-1 with −52.469 Kcal/mol, −56.043 Kcal/mol and −52.819 Kcal/mol free binding energies whereas Fostamatinib (control) showed slightly positive score −53.750 Kcal/mol. These three ligand molecules for each receptor were then subjected to induced fit docking (IFD) which is more powerful tool to predict the ligand-receptor interaction with better accuracy in pose prediction. One best ligand was selected for each ligand based on IFD score and those ligands were then analyzed for their drug potential.

#### 3.2.1. Binding Mode of 6-Gingerol with COX-2

6-Gingerol docked with COX-2 with an IFD score of −2474.460 Kcal/mol and XP Gscore of −9.264 Kcal/mol (**Table 5**). It formed 5 conventional hydrogen bonds with Gln454, His214, Tyr385, Thr212 and Asn382 amino acid residues inside the binding pocket of COX-2 at 2.01 Å, 2.29 Å, 1.93 Å, 3.60 Å and 2.52 Å respectively. It also formed 1 non-conventional hydrogen bond with His388 amino acids and few other hydrophobic interactions i.e., Pi-Pi Stacked, Pi-Pi T Shaped and Pi-Alkyl interactions with interacting amino acids inside the binding cleft of COX-2. It interacted with 11 amino acids in total inside the binding site of COX-2 (**Figure 4**).

#### 3.2.2. Binding Mode of Yakuchinone A with IKK

Yakuchinone A docked with IKK with an IFD score of −1081.370 Kcal/mol and XP Gscore of −9.111 Kcal/mol (**Table 5**). It formed 2 conventional hydrogen bonds with Lys106 amino acid residue inside the binding pocket of IKK at 5.46 Å and 2.00 Å distance apart respectively. Again, it formed 2 additional conventional hydrogen bonds with Glu100 and Cys99 amino acids at 2.72 Å and 1.96 Å distance apart respectively. Yakuchinone A formed 1 nonconventional hydrogen bond with Glu100 amino acid residue and few other Pi-Alkyl interactions with interacting amino acids inside the binding cleft of IKK. It interacted with 7 amino acids in total inside the binding site of IKK (**Figure 4**).

#### 3.2.3. Binding Mode of Yakuchinone B with TBK-1

Yakuchinone B docked with TBK-1 with an IFD score of −1309.340 Kcal/mol and XP Gscore of −7.971 Kcal/mol (**Table 5**). It formed 2 conventional hydrogen bonds with Ser93 and Cys89 amino acid residues inside the binding pocket of TBK-1 at 5.46 Å and 2.00 Å distance apart respectively. Again, it formed 3 additional non-conventional hydrogen bonds with Gly139 and Phe88 amino acids. Yakuchinone B formed few other hydrophobic interactions i.e., Pi-Sigma, Pi-Alkyl and Amide-Pi Stacked with interacting amino acids inside the binding cleft of TBK-1. It interacted with 7 amino acids in total inside the binding site of TBK-1 (**Figure 4**)

### 3.3. ADME/T Prediction

Best three selected ligand molecules (**Table 6**) were analyzed for their potential ADME/T profiles (**Table 7**). All of the ligand molecules were predicted to have high intestinal absorption and low oral bioavailability. Only Yakuchinone B has lower Caco-2 permeability. All of them are non-substrates of membrane P-glycoproteins and capable of penetrating blood brain barrier.

Only 6-Gingerol was reported to be inhibitor of P-glycoproteins. All of them were reported to be the substrate of CYP3A4 and 6-Gingerol and Yakuchinone A are substrates of CYP2D6. None of them showed sign of inhibition towards CYP3A4, CYP2D6 and CYP2C9.

None of the ligands was reported to be OCT2 (Organic Cation Transporter 2) substrate and show AMES toxicity and Hepatotoxicity. All of the ligands were reported to be inhibitors of *hERG* (Human ether-a-go-go related gene) channel. Only 6-Gingerol showed sign of eye irritation. All ligands showed Type III acute oral toxicity.

### 3.4. Pharmacological and Biological Activity Prediction

Best three ligand molecules were analyzed for their tentative pharmacological activities (**Table 8**). They were analyzed to understand their association with anti-inflammatory and other activities with enzymes, signaling proteins, transcription factors and cytokines involved in inflammatory cascades (**Table 8**).

Probability scores of intended pharmacological activities of investigated ligands varied with variety of extent and Yakuchinone B performed slightly better in the pharmacological activity prediction experiment followed by Yakuchinone B and 6-Gingerol. Yakuchinone B showed activities as TNF expression inhibitor and JAK2 expression inhibitor with probability score greater than 0.7. 6-Gingerol showed Macrophage colony stimulating factor agonist activity with probability score greater than 0.7. Other scores of intended activities by selected ligand molecules ranged from moderate to low.

Thereafter, the selected ligands were investigated for their biological activities against GPCR ligand, ion channels, enzyme etc. (**Table 9**). 6-gingerol showed better positive scores as Enzyme inhibitor, Nuclear receptor ligand and GPCR ligand modulator with higher positive probability scores. However, Yakuchinone A and Yakuchinone B also showed similar activities against few parameters.

**Table 9:** Result of biological activity prediction of best ligands.

| Bioactivity | Score |  |  |
| --- | --- | --- | --- |
|  | 6- Gingerol | Yakuchinone A | Yakuchinone B |
| GPCR ligand | 0.16 | 0.07 | -0.00 |
| Ion channel modulator | 0.04 | -0.02 | -0.17 |
| Kinase inhibitor | -0.33 | -0.31 | -0.36 |
| Nuclear receptor ligand | 0.20 | 0.12 | 0.17 |
| Protease inhibitor | 0.15 | 0.01 | -0.09 |
| Enzyme inhibitor | 0.38 | 0.16 | 0.13 |

**Table 10:** Result of DFT calculation. The unit of HOMO, LUMO, gap, hardness and softness are in Hartree and the unit of dipole moment is in Debye.

| Compound Name | HOMO | LUMO | Gap | Hardness ($\eta$) | Softness (S) | Dipole Moment |
| --- | --- | --- | --- | --- | --- | --- |
| 6-gingerol | -0.19924 | -0.01477 | 0.18447 | 0.09223 | 10.84167 | 3.3525 |
| Yakuchinone A | -0.19952 | -0.01594 | 0.18358 | 0.09179 | 10.89443 | 2.1614 |
| Yakuchinone B | -0.21682 | -0.05847 | 0.15835 | 0.07917 | 12.63104 | 4.5276 |

### 3.5. P450 Site of Metabolism Prediction

Best selected ligand molecules were examined for their potential sites of metabolism against three major isoforms of Cytochrome P450 family of enzymes i.e., CYP3A4; CYP2D6 and CYP2C9 (**Figure 5**). All of the selected ligand molecules were reported to have multiple atoms which are vulnerable to a specific enzyme of CYP450 family. 6-Gingerol showed almost similar sites of metabolism for all three isoforms but Yaluchinone A and Yakuchinone B showed different potential metabolism sites for metabolism by different emzymes.

### 3.6. Analysis of Frontier’s Orbitals

Detailed HOMO energy, LUMO energy, energy gap (HOMO-LUMO gap), hardness and softness of the selected three compounds are summarized in **Table 10** and the HOMO and LUMO occupation of the ligands is illustrated in **Figure 6** for each compound. Highest gap was observed for 6-Gingerol and lowest gap was observed for Yakuchinone B.

According to the energy gap the order of the compounds is: 6-Gingerol >Yakuchinone A> Yakuchinone B. Along with the HOMO and LUMO energy, Dipole moment of the selected ligand molecules were also calculated and according to dipole moment the order of compoiunds: Yakuchinone A > 6-Gingerol > Yakuchinone B.

## 4. Discussions

*In silico* analysis of drug likeness helps to filter out compounds with poor drug like potentials usually those have poor physicochemical properties. Violation of Lipinski’s rule by a compound indicates that the compound is more likely to fail in the later trial [84][85]. In this experiment the ligand molecules were analyzed in accordance with Lipinski’s rule of five. Cassumunin A and Cassumunin B violated the standard rule and then they were removed from consideration of further experiment (**Table 3**). Thereafter, 12 ligand molecules except these two were analyzed in the molecular docking experiment. Molecular docking is one of most commonly used tools in computer-aided drug designing. This tool works on specific algorithm which assigns binding energy after docking which in turn reflects binding affinity of a ligand to a molecular target [56][86][87]. Lowest binding energy of ligand-receptor complex reflects higher affinity meaning they remain more time in contact [88]. In this experiment SP and XP ligand docking were carried out to make comparison among docking parameters of different ligands. However, best three ligands for one receptor were selected based on MM-GBSA scoring because it is more rigorous scoring method (**Table 4**) [89][90]. Selected three ligands showed better binding free energies than approved inhibitors (control). Then the selected ligand molecules were subjected to induced fit docking (IFD) which is even powerful tool to generate poses and assigning scores [91]. Upon continuous exploration with different methods of molecular docking, 6-Gingerol, Yakuchinone A and Yakucinone B were selected as the best inhibitor of COX-2, IKK and TBK-1 respectively (**Table 5** & **6**). Hydrogen bonding and hydrophobic interactions play significant role in strengthening the ligand receptor interactions [92]. All selected ligands were reported to form multiple hydrogen bonds and other forms of hydrophobic interactions inside the binding cleft of respective receptors (**Figure 4**).

*In silico* analysis of absorption, distribution, metabolism, excretion and toxicity is crucial to determine whether a drug is likely to survive in the later stages of drug development and these data again help to reduce the time and cost of drug discovery approach by assisting *in vitro* assays [71][93]. Blood brain barrier permeability is a major concern for the drugs targeting primarily the cells of central nervous system (CNS). Oral delivery system is the most commonly used route of drug delivery and the delivered drug migrates through the digestive tract into the intestine and so the drug under investigation is appreciated to be highly absorbed in human intestinal tissue. P-glycoproteins are the cell membrane glycoproteins that are responsible to facilitate the transport of many drugs through the cell membrane and hence their inhibition by candidate drugs may affect the normal drug transport inside human body. *Caco2* permeability to drug reflects the human intestinal tissue permeability since this cell line is commonly used for *in vitro* permeability assay [94]-[96]. Cytochrome P450 family of enzymes is center to control the drug interaction, metabolism inside human body and excretion outside the body. Inhibition of these enzymes may lead to acute drug toxicity, slow clearance and eventually malfunction of the drug compound inside human body [97]-[99]. AMES toxicity approach is used to examine the toxicity endpoint of chemicals in question [100][101]. *hERG* (*Human ether-a-go-go related gene*) channels are the voltage gated potassium ion channels that play key roles for potassium ion transport through the cell membrane. Different structurally and functionally unrelated drugs have been reported to block the *hERG* potassium channel which has raised the concern of off-target drug interaction and so, screening compounds for activity on *hERG* channels early in the lead optimization process is crucial [102]. Renal OCT2 (organic cation transporter 2) is important for drug and xenobiotic excretion through kidney. The substrates of this transporter protein are considered to easily be excreted with urine [103]. All of the selected ligands exhibited almost similar properties in ADME/T test (**Table 7**).

Pharmacological activity (PASS prediction) is predicted in the context of probability of activity (Pa) and Probability of inactivity (Pi) of a compound and the result of the prediction varies between 0.000 and 1.000. The activity is considered possible When Pa>Pi [104]. When Pa>0.7, the compound is very likely to exhibit the activity but possibility of the compound being analogue to a known pharmaceutical is also high. Compound with 0.5< Pa <0.7 score is likely to exhibit the activity but the probability is less along with the chance of being a known pharmaceutical agent is also lower. When Pa<0.5, then the compound is less likely to exhibit the activity [105]. Pharmacological activity was predicted for the compounds against anti-inflammatory activity and other proteins, transcription factors, enzymes and cytokines involved in different inflammatory cascades. Yakuchinone B performed slightly better almost overall all the ligands were predicted to have similar activities (**Table 8**). Then the ligands were analyzed for their potential biological activities against GPCRs (G protein coupled receptors), ion channels, enzymes, nuclear receptors etc. which are the most potent drug targets in human body. Only GPCRS are the targets of 50% of currently available drugs in the market [106]-[108]. The ligands showed significant connections (probability scores) against all targets which might be useful for drug discovery expenditure but at the same time could also raise concern of unpleasant drug-target interaction as useless (**Table 9**).

Then the ligands were analyzed for their potential metabolism sites for three major enzymes i.e., CYP3A4, CYP2D6 and CYP2C9 of Cytochrome P450 family and multiple sites for each molecules were recorded (**Figure 5**). The best ligands were also analyzed for their HOMO and LUMO energy and occupation. HOMO is usually a constraint portion in a molecule that is capable of donating electrons whereas LUMO is responsible for accepting electrons (**Figure 6**). HOMO-LUMO gap is used to define the stability of a compound. A compound having highest gap is likely to undergo a chemical reaction more easily [109][110]. Yakuchinone A and 6-Gingerol showed almost similar energy gap whereas Yakuchinone B showed slightly lower gap (**Table 10**).

Finally, 6-Gingerol, Yakuchinone A and Yakuchinone B were the best findings of this study. Anti-inflammatory activities of these compounds have already been proven in laboratory experiments [111]-[113]. These compounds also performed quite similar in different post-screening study after docking experiment which could be useful for further drug discovery approach. However, these findings might be required to be supported by further in vitro study.

Overall, this study recommends 6-Gingerol, Yakuchinone A and Yakuchinone B as the best inhibitors of COX-2, IKK and TBK-1 respectively among the selected curcumin analogues although other compounds can also be investigated since they also performed well in docking experiment.

## 5. Conclusion

In this study 14 Curcumin analogues were utilized to explore their anti-inflammatory activities against three signaling proteins in the TLR4 pathway. Upon continuous computational exploration 6-Gingerol, Yakuchinone A and Yakuchinone B were identified as the best agents having inhibitory effects on COX-2, IKK and TBK-1 respectively. Their drug like potentials were also analyzed and they were found to have sound and similar drug like parameters. Therefore, these compounds could be considered as potential anti-inflammatory agents in the search for new medication against inflammation. However, authors suggest further *in vitro* study with these compounds to confirm their anti-inflammatory activities and strengthen these findings.

## Data Availability Statement

Authors made all the data generated during experiment and analysis available within the manuscript.

## Acknowledgements

Authors acknowledge the members of Swift Integrity Computational Lab, Dhaka, Bangladesh, a virtual platform of young researchers for their support during the preparation of the manuscript.

